# Genetic structure of a patchily distributed philopatric migrant: implications for management and conservation

**DOI:** 10.1101/216069

**Authors:** Anastasios Bounas, Dimitris Tsaparis, Ron Efrat, Gradimir Gradev, Marco Gustin, Kresimir Mikulic, Airam Rodríguez, Maurizio Sarà, Georgios Kotoulas, Konstantinos Sotiropoulos

## Abstract

Significant demographic fluctuations can have major genetic consequences in wild populations. The Lesser Kestrel (*Falco naumanni*) has suffered from both population declines and range fragmentation during the second half of the 20th century. In this study we analysed multilocus microsatellite data to assess the genetic structure of the species. Our analysis revealed significant genetic structuring of Lesser Kestrel populations, not only at cross-continental scale, but also regionally within Central and Eastern (CE) Mediterranean. We detected signs of genetic bottlenecks in some of the peripheral populations coupled with small effective population sizes. Values of genetic differentiation among the largest populations were low, albeit significant, whereas the small peripheral CE Mediterranean populations showed higher levels of differentiation from all other populations. Gene flow levels were relatively low among the discontinuously distributed populations of the CE Mediterranean region. We argue that the observed spatial genetic structure can be attributed at some level to the past demographic decline experienced by the species. Finally, we identify management units in the region, and inform the design of conservation actions aiming at the increase of population sizes and dispersal rates among peripheral populations.

## INTRODUCTION

In many animal species, the patterns of genetic differentiation and gene flow are highly influenced by the geographical characteristics of their habitats as well as their migratory behavior (Willoughby *et al*., 2017). Birds, and specifically raptors, can show long-distance migratory behavior and also exhibit natal and breeding site fidelity. As a general pattern, migratory populations of raptors have been found to have weaker genetic structure and higher genetic diversity than resident populations (Miller *et al*., 2012; Willoughby *et al*., 2017). For example some individuals might migrate through a different route upon their return on the breeding grounds, and consequently disperse and breed far from their natal site (Garcia *et al*., 2011). On the other hand, philopatry reduces or even inhibits gene flow among populations leading to increased genetic differentiation. Patchily distributed and locally isolated populations are susceptible to a greater influence of genetic drift which may result in a decrease in genetic diversity and fitness, thus compromising a species’ ability to adapt to a changing environment (Amos & Balmford, 2001). Genetic drift and inbreeding are expected to be stronger in peripheral populations relative to core ones, due to their small population sizes and low immigration rates (Vucetich & Waite, 2003; Hanski & Gaggiotti, 2004). Therefore, the assessment of genetic structure and the identification of its underlying processes become essential tasks providing valuable information towards the design and implementation of effective conservation strategies. For example, identification of management units (MUs, Moritz, 1994; Taylor & Dizon, 1999; Palsbøll, Berube & Allendorf, 2007) would be central to delineate populations for monitoring and thus aid their short-term management.

The Lesser Kestrel (*Falco naumanni*, Fleischer, 1818) is a small migratory falcon breeding from the Mediterranean basin across Middle East and Central Asia to Mongolia and China, and wintering in sub-Saharan Africa (Cramp & Simmons, 1980). The species underwent rapid declines throughout its European range in the early 1960s mainly as a result of agricultural intensification and subsequent habitat degradation and land use changes (Iñigo & Barov, 2010). In Central-Eastern Mediterranean region (hereafter CE Mediterranean) and especially in the Balkan Peninsula, the decline was dramatic, leading to local extinctions and consequently to significant range contraction and fragmentation (Iñigo & Barov, 2010). Currently, the species shows a patchy distribution and is considered to have a “depleted” status in the region (BirdLife International, 2017a). The two largest (core) populations are located in the Apulia-Basilicata area of southern Italy and in Central Greece, still holding several thousand pairs (BirdLife International, 2017a). Peripheral populations of smaller size, still exist in the Former Yugoslav Republic of Macedonia (Uzunova & Lisichanets, 2016), in Sicily (Sarà, 2004), the European part of Turkey (Kmetova *et al*., 2012) and throughout continental Greece as well as on some of the Greek islands (Legakis & Maragou, 2009). Finally, a small geographically isolated population is located in Croatia, at the northernmost edge of the species European distribution (Mikulic *et al*., 2013). Certain conservation actions, mainly implemented in Western Europe, have led to a stabilization and slightly positive population trend and subsequently the down-listing of the species from Vulnerable to Least Concern (BirdLife International, 2017b).

Several previous studies have examined the genetic structure of the species and the underlying processes, at both continental (Wink, Sauer-Gurth & Pepler, 2004; Alcaide *et al*., 2008a; Alcaide *et al*., 2008b) and finer spatial scales (Ortego *et al*., 2008a; Ortego *et al*., 2008b; Alcaide *et al*., 2009; Di Maggio *et al*., 2014). It has been proposed that Asian Lesser Kestrels are considerably differentiated from European conspecifics, whereas populations across the Western Palearctic seem to follow an isolation by distance pattern while maintaining high levels of genetic diversity (Wink *et al*., 2004; Alcaide *et al*., 2008b). On a regional scale though, it seems that the species’ philopatry does not lead to fine-scale genetic structuring (Alcaide *et al*., 2009); nevertheless, population structure could emerge, depending on the size and the degree of spatial isolation and the levels of gene flow among populations (Ortego *et al*., 2008b). Such restricted gene flow patterns do not necessarily result from physical barriers hindering dispersal, especially in efficient dispersers such as the Lesser Kestrel, but can be attributed also to their philopatric behaviour. Indeed, the species shows high philopatry, with adult birds returning to breed close to their previous breeding territories (Negro, Hiraldo & Donázar, 1997), however, juveniles disperse more and in greater distances (Serrano & Tella, 2003; Bounas *et al*., 2016a). At least at a continental scale, Lesser Kestrels show strong migratory connectivity, i.e. there is a spatial segregation of breeding populations at the wintering range: European populations winter in the Sahel, while Asian populations winter in east and South Africa (Wink *et al*., 2004; Rodriguez *et al*., 2009)

Herein, we examine the genetic structure of the Lesser Kestrel, 1) across the broader species range and 2) within the CE Mediterranean. We aim to identify patterns of genetic variation and gene flow among populations as well as their underlying processes. Such information could be of crucial importance for conservation programs to identify the need of local scale conservation actions and inform their design.

## MATERIAL AND METHODS

### Population sampling and DNA extraction

Samples were obtained from 12 breeding sites in CE Mediterranean, where the species shows a fragmented distribution as well as from Mongolia, Israel and two sites from Spain. Individuals sampled in Bulgaria originated from Spain and were used for the reinforcement of the species, thus treated as if they were sampled in Extremadura (ES/BG; Fig. 1). A total of 295 individuals were sampled during four consecutive breeding seasons (2013 – 2016) with the exception of Spanish samples (SES) that were collected in 2007. Birds were caught by hand in the nest or using mist nets or spring traps close to the nest. To minimize biases associated with relatedness only a single fledgling per brood was sampled. Two drops of blood (≈50μl) were obtained from each individual by leg-pricking and immediately stored in blood storage cards (NucleoCards^®^) at room temperature until DNA extraction. DNA was extracted using the NucleoSpin Tissue kit (Macherey-Nagel) following the manufacturer’s protocol.

**Figure 1.**
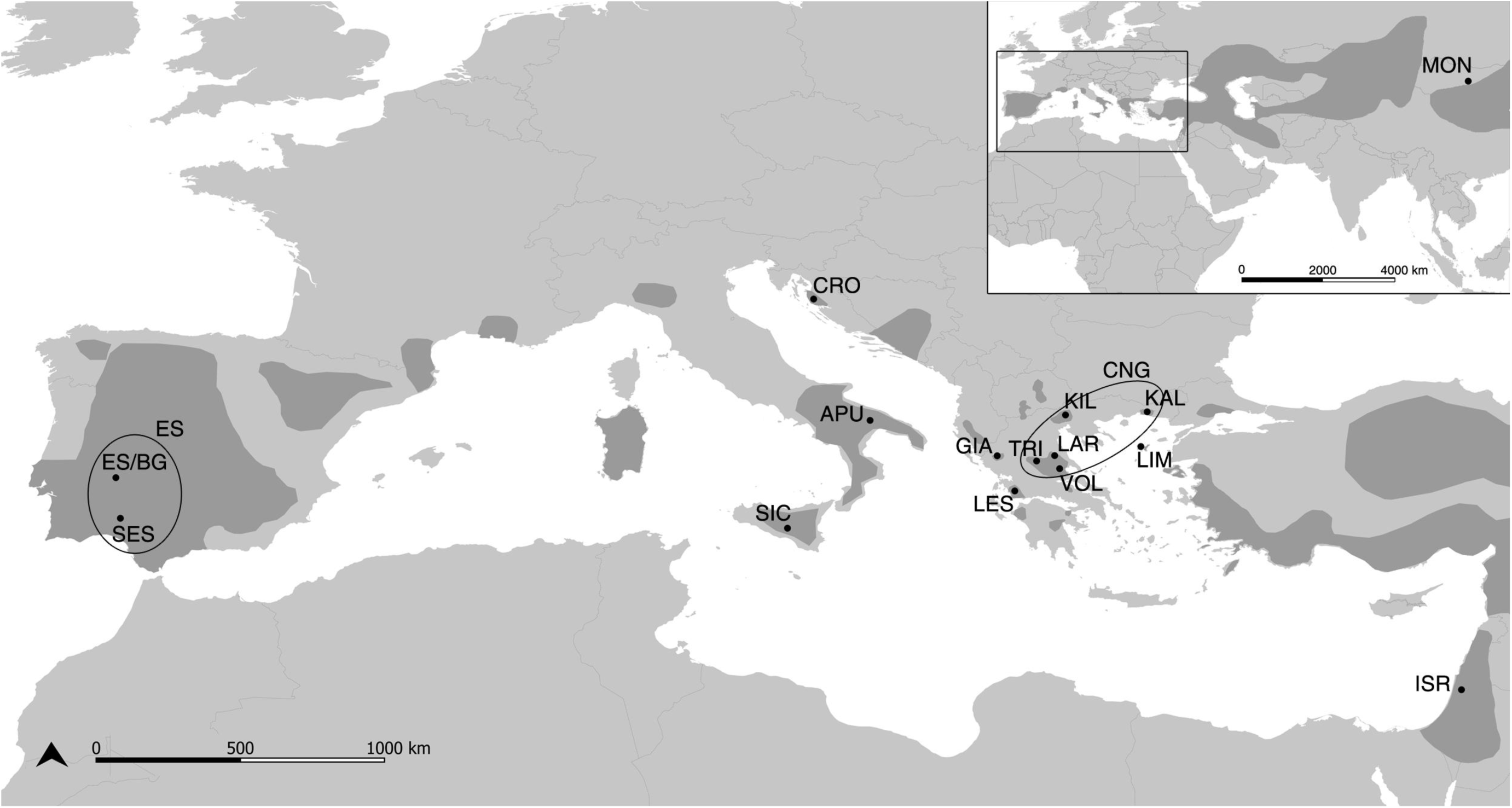
Map of the Lesser Kestrel populations used in the present study. SES: Andalucia, ES/BG: Extremadura, APU: Apulia, SIC: Sicily, CRO: Croatia, GIA: Ioannina, LES: Agrinio, TRI: Trikala, LAR: Larisa, VOL: Volos, KIL: Kilkis, KAL: Komotini, LIM: Limnos, ISR: Israel, MON: Mongolia. Shaded areas represent the breeding distribution of the Lesser Kestrel (modified from BirdLife International). Sampling sites pooled for the analyses are circled (ES: Spain; CNG: Central-North Greece)

### Microsatellite amplification and genotyping

Each individual was genotyped at a total of 18 microsatellite loci. Seven loci were originally isolated from the Peregrine Falcon *Falco peregrinus* (Nesje *et al*., 2000; Alcaide *et al*., 2008a), whereas 11 were developed specifically for the Lesser Kestrel (Ortego *et al*., 2007; Padilla *et al*., 2009). Details on loci properties and primers used for their amplification are presented in Supporting information, Table S1. All loci were amplified in five multiplex reactions using forward 5’-fluorescent-labelled primers and the KAPA2G Fast Multiplex PCR Kit (Kapa Biosystems). Each 12.5 μl reaction contained 2pM of each primer and 1x KAPA2G Mix and was carried out using the following profile: an initial denaturation step of 3 min at 95 °C, 30 cycles of 15 s at 95°C, 30 s at 60 °C, 30 s at 72 °C, with a final extension step of 10 min at 72 °C. PCR products were separated and visualized using an ABI 3730xl capillary sequencer (Applied Biosystems) and genotypes were scored by eye with STRand v.2.4.59 (Toonen & Hughes, 2001). Randomization of samples was employed throughout lab processes to avoid any plate/gel specific errors that might lead in population specific biases (Meirmans, 2015). In addition, a subset of 50 individuals was re-genotyped to quantify error rates due to allelic dropout or genotyping errors but no inconsistencies were detected. We used the package “MsatAllele” (Alberto, 2009) in R 3.2.2 (R Core Team, 2015) to allocate alleles to their respective size classes. Genotyping errors, due to null alleles and stuttering, were examined for all loci and sampled populations using MICROCHECKER (Van Oosterhout *et al*., 2004). Two loci (Fnd1.2 and Fnd2.1) were omitted from further analyses due to the presence of null alleles and extensive stuttering respectively, and thus the full analysis was based on the remaining 16 microsatellite loci.

### Genetic analyses

To evaluate the genetic variability in each population, standard genetic diversity indices (*A*: number of alleles, *H*_O_: observed and *H*_E_: expected heterozygosity) were calculated using the program GENALEX v.6.5 (Peakall & Smouse, 2012). Rarefied Private Allelic richness (*π*) estimates were produced using HP-RARE (Kalinowski, 2005). Allelic richness (*A*_R_) corrected for different sample sizes was calculated using FSTAT 2.9.3.2 (Goudet, 2002). The same software was used to calculate the inbreeding coefficient (*F*_IS_), test for deviations from Hardy-Weinberg proportions at each locus and sampled population, as well as for linkage disequilibrium (LD) using 1000 randomizations and adjusting significance for multiple comparisons (adjusted P value < 0.0003). In addition we performed a Chi-Square test and Fisher’s method to confirm the Hardy Weinberg (HW) equilibrium results.

The software Arlequin 3.5.1.3 (Excoffier & Lischer, 2010) was used to calculate F_ST_ values between all pairs of sampled populations and test them for statistical significance using 10000 permutations. In addition to F_ST_, we also calculated Jost’s D (D_est_) as an unbiased estimator of differentiation that performs better than other relatives, in cases of markers with different number of alleles (Gerlach *et al*., 2010) as is the case in this study (see Supporting information, Table S1). Pairwise D values (mean D_est_) among populations were calculated with the R-package “DEMEtics” (Gerlach *et al*., 2010) and statistical significance was tested using 1,000 bootstrap iterations. For both estimators, p-values were adjusted for multiple comparisons after the B-H method (Benjamini & Hochberg, 1995). We used IBDWS v.3.23 (Jensen, Bohonak & Kelley, 2005) in order to obtain any statistically significant associations between pairwise genetic (F_ST_/[1-F_ST_]) and linear geographic (log km) distance matrices using 30,000 randomizations. The analysis was performed both on the full and the reduced (i.e. CE Mediterranean) datasets.

To evaluate the genetic population structure, the Bayesian clustering software STRUCTURE 2.3.4 (Pritchard, Stephens & Donnely, 2000) was used to infer the number of genetically homogeneous clusters present in the dataset. The analysis was conducted both on the full dataset as well as only on the populations of CE Mediterranean region. For both analyses we assumed the admixture ancestry model and correlated allele frequencies (Falush, Stephens & Pritchard, 2003), using sampling location as prior information (LOCPRIOR model; Hubisz *et al*., 2009) as it is deemed to be more sensitive at inferring population structure at lower levels of divergence, which is expected in good dispersers such as birds. Runs were set with a burn-in period of 2 × 10^5^ iterations followed by 10^6^ MCMC steps with 20 replicates for each K value (1 to 15 for the full dataset and 1 to 12 for CE Mediterranean). STRUCTURE runs were implemented on a beowulf cluster using PARASTRUCTURE Perl script (Lagnel, 2015) and STRUCTURE plots were constructed using DISTRUCT (Rosenberg, 2004). The most likely value of genetic clusters, K, was evaluated following the ΔK method (Evanno, Regnaut & Goudet, 2005) implemented in STRUCTURE HARVESTER (Earl & Vonholdt, 2012), as well as by calculating the posterior probability for each K. We present all cases with high probability that warrant a biological interpretation (Meirmans, 2015) and are supported by both STRUCTURE runs.

Finally, the breeding populations of CE Mediterranean were grouped according to Structure results as well as F_st_ and D_est_ calculations, and the directional contemporary gene flow and its relative magnitude among them was estimated using the divMigrate function (Sundqvist et al. 2016) in the R-package “diveRsity” (Keenan et al. 2013). The method provides a relative (to within the analysis) migration network graph aiming to visualize the gene flow patterns among populations, with the resulting metric representing a proportion of the gene flow among areas, scaled to the largest magnitude estimated. Populations are represented as nodes and the properties of the lines connecting them are based on the relative strength of gene flow. Populations that exhibit strong gene exchange between them but weak gene flow with others tend to cluster closely together, reflecting patterns of genetic structure. The method is described in detail in Sundqvist *et al*. (2016). We used *N*_M_ (Alcala, Goudet & Vuilleumier, 2014) as a measure of genetic distance, and tested whether gene flow among populations was asymmetrical using 10000 bootstrap iterations.

Signs of bottlenecks were evaluated in grouped populations using three approaches. First we calculated *M*, a ratio based on the number of alleles to the allelic size range (Garza & Williamson, 2001) in Arlequin. M will be smaller in populations that have suffered a decline than in populations that are in mutation-drift equilibrium. A test for heterozygosity excess was performed in BOTTLENECK (Piry, Luikart & Cornuet, 1999) using the Wilcoxon signed rank test running 1000 iterations and using the two-phase model (TPM). As the microsatellites we used are either of dinucleotide perfect repeats or of imperfect repeats, both of which may tend toward the infinite allele model (IAM; Cornuet and Luikart, 1996), we fixed the proportions of the TPM in favour of the IAM (Cristescu *et al*., 2010) including 20% of the stepwise mutation model (SMM) and 80% of IAM. We finally tested for a mode-shift distortion using a graphical approach, by plotting the number of alleles in 10 allele frequency classes with 0.1 intervals, according to Luikart *et al*. (1998). In a population at equilibrium, alleles with frequencies in the first class (<0.1) are expected to be more numerous than those belonging to the second class, therefore allele frequencies present a characteristic L-shaped distribution. Plots were examined for mode-shift distortions that would be consistent with a bottlenecked population (Luikart *et al*., 1998; Cristescu *et al*., 2010). Plots were produced in R 3.2.2 and allele frequency calculations were performed using the R package “Gstudio” (Dyer, 2014). Finally, the effective population size (*N*_E_) along with 95% confidence intervals (CIs) were estimated using the bias-corrected version of the method based on linkage disequilibrium (Hill, 1981; Waples, 2006) as implemented in NeESTIMATOR v.2.01 (Do *et al*., 2014). N_E_ estimates were obtained for all but the core populations (ES, APU, CNG) and Mongolia, as it is very difficult to obtain reliable estimates for large populations using this method (Waples & Do, 2010).

## RESULTS

All microsatellite markers were found to be polymorphic across populations and the number of alleles per locus ranged from four (loci Fnd1.4, Fp86-2, Fp89) to 38 (locus Fnd1.6; Supporting information, Table S1).

### Genetic diversity

We did not detect any differences in the mean allelic richness across populations. The average *H*_O_ was 0.64, *H*_O_ and *H*_E_ were similar and *F*_IS_ was not significant in all cases (Table 1). Private alleles were present in all sampled populations with the exception of Croatia (CRO), Komotini (KAL) and one subpopulation in Central Greece (TRI). Generally, populations did not deviate from Hardy-Weinberg proportions but some deviations of individual loci were detected: four populations (CRO, LIM, ISR, MON) showed deviations at two, one, one and two out of the 16 loci respectively (see Supporting information, Table S4). Since these loci did not show consistent deviations across all populations, we included them in subsequent analyses attributing this disequilibrium in processes specific to those populations. No LD was detected between any of the loci across all populations.

**Table 1.** Measures of genetic variation of all sampled Lesser Kestrel populations. Number of genotyped individuals (*N*), number of alleles per locus (*A*), allelic richness (*A*_R_), observed (*H*_O_) and expected (*H*_E_) heterozygosity, private allelic richness (*π*) and inbreeding coefficient (*F*_IS_). Values are presented as means ± standard error (in brackets)

| Code | Location | $N$ | $A$ | $A_R$ | $H_O$ | $H_E$ | $\pi$ | $F_{IS}$ |
| --- | --- | --- | --- | --- | --- | --- | --- | --- |
| SES | Andalucia | 19 | 7.1<br>(0.9) | 5.4<br>(0.6) | 0.635<br>(0.037) | 0.696<br>(0.037) | 0.17 | 0.082<br>(0.041) |
| ES/BG | Extremadura | 25 | 8.3<br>(1.3) | 5.6<br>(0.7) | 0.674<br>(0.040) | 0.699<br>(0.037) | 0.26 | 0.029<br>(0.037) |
| APU | Apulia | 44 | 9.4<br>(1.5) | 5.5<br>(0.6) | 0.669<br>(0.038) | 0.707<br>(0.036) | 0.13 | 0.053<br>(0.025) |
| SIC | Sicily | 12 | 5.9<br>(0.8) | 5.1<br>(0.6) | 0.682<br>(0.038) | 0.667<br>(0.041) | 0.18 | -0.045<br>(0.045) |
| CRO | Croatia | 14 | 5.6<br>(0.6) | 4.7<br>(0.5) | 0.654<br>(0.048) | 0.650<br>(0.048) | 0.03 | -0.023<br>(0.043) |
| GIA | Ioannina | 24 | 7.6<br>(1.1) | 5.2<br>(0.6) | 0.634<br>(0.051) | 0.654<br>(0.048) | 0.16 | 0.036<br>(0.031) |
| LES | Agrinio | 16 | 6.7<br>(0.8) | 5.4<br>(0.6) | 0.603<br>(0.058) | 0.684<br>(0.042) | 0.18 | 0.136<br>(0.047) |
| TRI | Trikala | 20 | 7.6<br>(1.3) | 5.5<br>(0.7) | 0.626<br>(0.056) | 0.680<br>(0.046) | 0.08 | 0.087<br>(0.046) |
| LAR | Larisa | 20 | 7.4<br>(1.1) | 5.4<br>(0.6) | 0.669<br>(0.053) | 0.680<br>(0.044) | 0.09 | 0.024<br>(0.045) |
| VOL | Volos | 20 | 7.6<br>(1.2) | 5.5<br>(0.7) | 0.678<br>(0.051) | 0.662<br>(0.048) | 0.13 | -0.027<br>(0.035) |
| KIL | Kilkis | 13 | 6.4<br>(0.9) | 5.3<br>(0.7) | 0.621<br>(0.062) | 0.661<br>(0.048) | 0.09 | 0.067<br>(0.064) |
| KAL | Komotini | 20 | 7.1<br>(1.1) | 5.3<br>(0.6) | 0.689<br>(0.053) | 0.677<br>(0.046) | 0.04 | -0.019<br>(0.040) |
| LIM | Limnos | 11 | 5.1<br>(0.6) | 4.8<br>(0.5) | 0.661<br>(0.060) | 0.667<br>(0.041) | 0.09 | 0.024<br>(0.060) |
| ISR | Israel | 20 | 7.3<br>(1.1) | 5.5<br>(0.7) | 0.619<br>(0.062) | 0.682<br>(0.042) | 0.28 | 0.124<br>(0.055) |
| MON | Mongolia | 17 | 7.2<br>(1.0) | 5.4<br>(0.6) | 0.585<br>(0.057) | 0.655<br>(0.049) | 0.52 | 0.155<br>(0.046) |

### Population structure

Pairwise F_ST_ and D_est_ values were highly correlated (Pearson’s correlation: *r* = 0.95, *P* < 0.001). Estimates of both F_ST_ and D_est_ (Table 2) varied between population pairs (F_ST_: 0.003 − 0.06; D_est_: 0.02 − 0.19), with all the core populations of Europe (Spain, Apulia, Central Greece) showing low differentiation among them. On the other hand, Mongolia and Israel seem differentiated from all others, while the small peripheral CE Mediterranean populations showed higher levels of differentiation from all other populations. This was particularly evident for the populations of Croatia and Limnos where the highest values of D_est_ were recorded when compared to all other populations (Table 2). Mantel tests showed that genetic distance is not significantly correlated with geographical distance across all sampled populations (*r* = 0.33, *P* = 0.1, Supporting information, Figure S1). Conversely, genetic divergence of CE Mediterranean populations correlated significantly with the geographical distance among them (*r* = 0.57, *P* = 0.01, Supporting information, Figure S1).

**Table 2.**
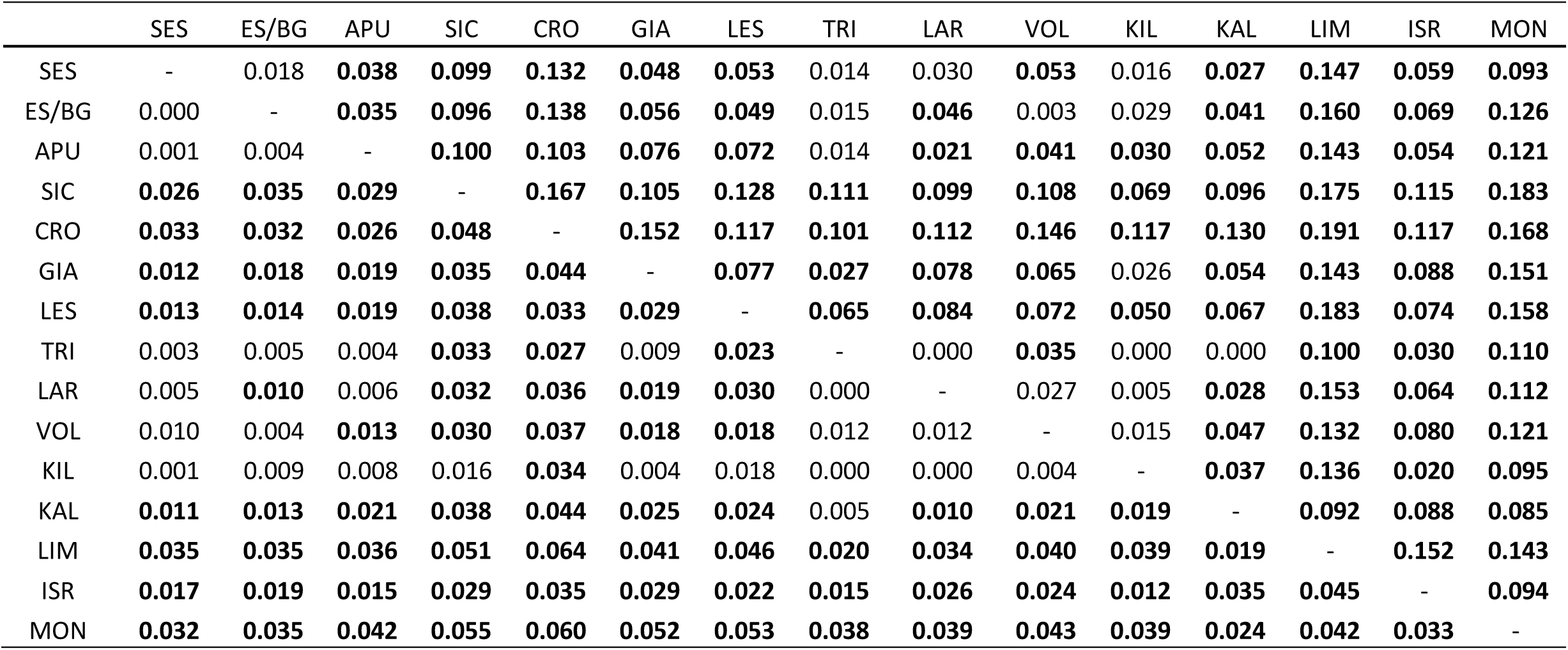
Pairwise F_st_-values (below diagonal) and D_est_-values (above diagonal) among Lesser Kestrel populations. Statistically significant values after B-H correction for multiple comparisons are given in bold (p < 0.039)

The Bayesian clustering method implemented in STRUCTURE suggested the presence of population structuring. When all individuals were modeled, the ΔK-method suggested two clusters (K = 2) as the most likely population structure (although the posterior probability was higher for K = 3; Supporting information, Figure S2, Table S2). For K = 2 the admixture model indicated two gene pools with all European populations showing high membership coefficients in the first cluster, and the MON and LIM populations belonging to the second (Fig. 2). This second gene pool seems to be present in all populations in Northern and Central Greece (CNG) whereas absent from all other western populations. Lesser Kestrels from Israel (ISR) were found to be highly admixed, exhibiting a mixed ancestry from both clusters (Fig. 2). For K = 3, the model adds another gene pool that is mostly represented by the Trans-Adriatic populations of Croatia (CRO), Apulia (APU) and Sicily (SIC) whereas Israel (ISR) still seems to be of mixed ancestry. All subpopulations within Spain (SES, ES/BG) and Central Greece (TRI, LAR, VOL) showed identical admixture proportions among them, implying no further substructure (Fig. 2).

**Figure 2.**
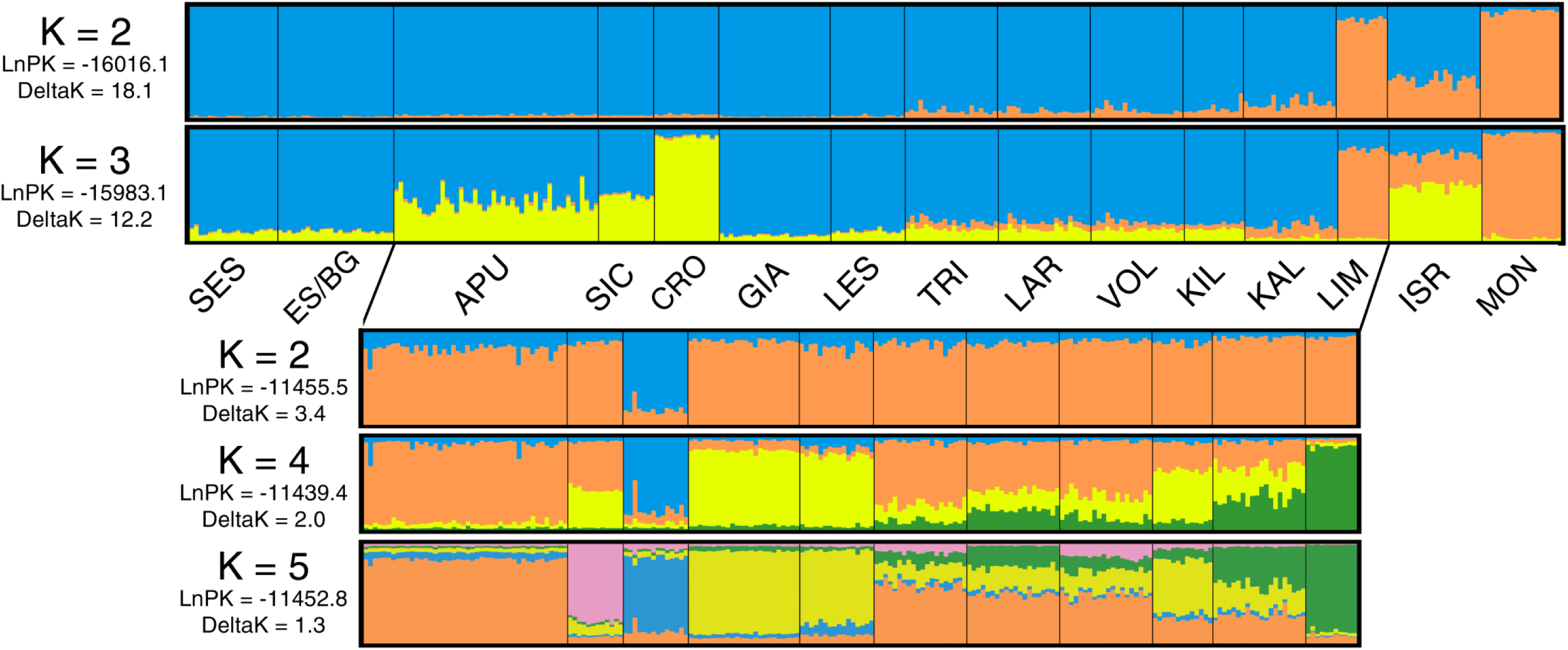
Admixture proportions (proportions of membership to each of K inferred clusters) of individual Lesser Kestrels. Upper plots correspond to the number of clusters when the full dataset was modeled (K = 2, K = 3), whereas the lower 3 plots show admixture proportions of the CE Mediterranean populations only, for K=2, K=4 and K=5 inferred clusters respectively

When individuals from the CE Mediterranean were modeled separately, the ΔK-method suggested K = 2 as the optimal number of clusters, whereas posterior probability estimates suggested the presence of four clusters (Supporting information, Figure S3, Table S3). The populations of Croatia (CRO) and Limnos (LIM) constitute two different genetic clusters, the one present in Limnos (LIM) extending throughout Greek populations from Northern to Central Greece (CNG), while two other clusters are present in all other populations in different proportions (GIA and LES populations of western Greece form a single group). The maximum value of five clusters further partitions the Sicilian population (SIC) as a separate group. It is apparent that there is no substructure in the Central and Northern Greek population, CNG (Fig. 2).

Because of the lack of any structure among the subpopulations of Central Greece (TRI, LAR, VOL) and Kilkis (KIL), we pooled the individuals from KIL into the central Greek group (CGR) to estimate the directional relative migration networks using divMigrate. The migration network (Fig. 3) reflects at some level the genetic structuring revealed by STRUCTURE analysis, as the core populations of the region (APU, CGR) cluster closely in the network space showing high gene flow and relatively low differentiation between them. The GIA and KAL populations also exhibited a relatively high gene flow with the core populations, with the former (GIA) showing connection with both APU and CGR while the latter (KAL) presents high gene flow rates only with CGR. The rest of the groups showed relatively reduced gene flow towards the core populations. It should be noted that as a general pattern, all peripheral populations seem to exchange migrants exclusively with the core populations but not between them, appearing isolated from each other (i.e. a star-shaped pattern of the migration network; Fig. 3), resembling a mainland-island metapopulation type (and not a patchy population type). However, there was no evidence of significantly asymmetric gene flow between any pair of populations.

**Figure 3.**
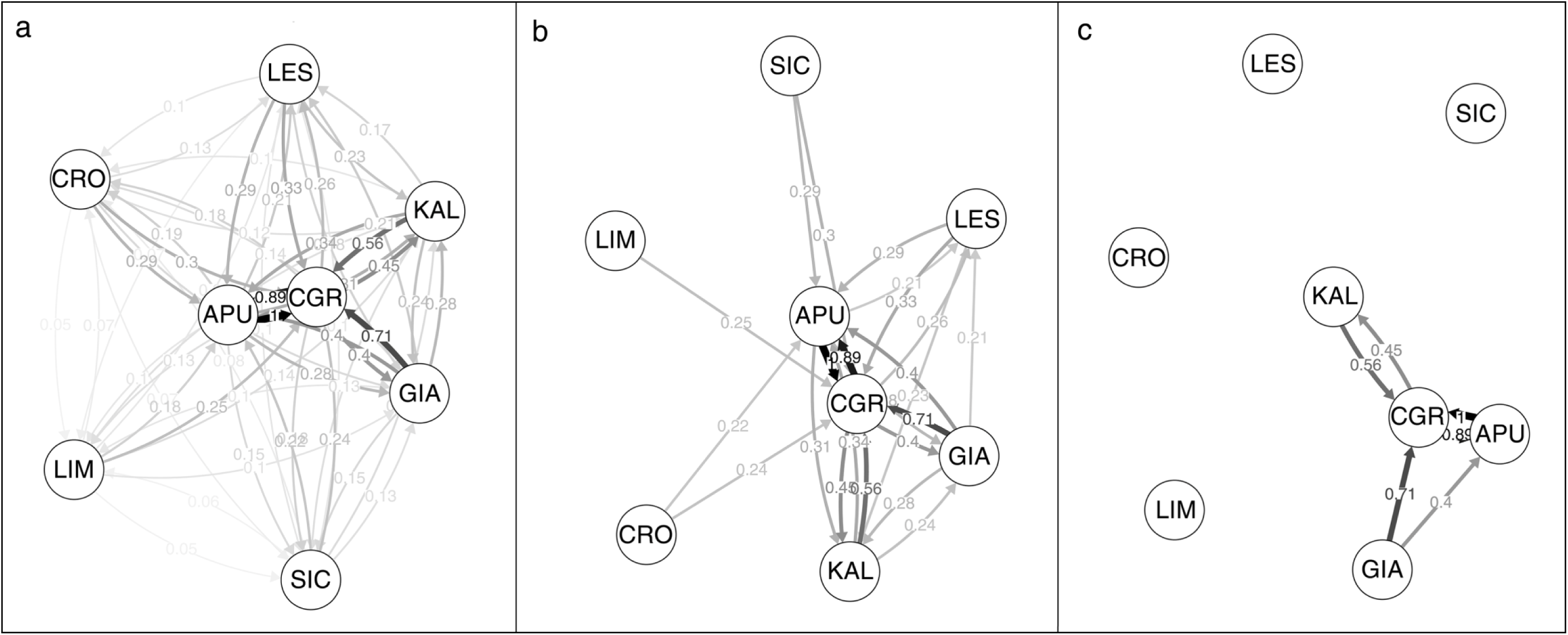
Directional relative migration networks of grouped Lesser Kestrel populations from CE Mediterranean. Networks were visualized with divMigrate using Alcala’s Nm. (a) all relative migration rate values (b) only values above 0.2 threshold (c) only gene flow values estimated >0.4. Line shading and thickness increases with the relative strength of gene flow.

### Demographic parameters

*M* ratios were found to be lower than the threshold value of 0.68 in all populations, which according to Garza & Williamson (2001) suggests that all populations have suffered a past bottleneck event. The Wilcoxon signed rank test conducted in BOTTLENECK software, detected signs of a recent population bottleneck for Sicily (SIC), Limnos island (LIM) and Israel (ISR) populations (Table 3). On the other hand, inspection of plots of allele frequency classes for recent bottlenecks did not reveal any mode-shift distortion in any of the populations. However, they did show that the Croatian population (CRO) is moving towards allele fixation, since this was the only population that exhibited an allele frequency in the class 0.9-1.0 (see Supporting information, Figure S4). Most of the peripheral populations were found to have small effective population sizes (Table 3), ranging from 24.2 (LIM) to 88.4 (LES). However, in some cases (SIC, LES and ISR populations) results should be treated with caution as 95% CIs were broad.

**Table 3.** Contemporary effective population sizes (*N*_E_) and 95% CI, results of tests for genetic bottlenecks: Garza-Williamson M values, Wilcoxon sign-rank tests for heterozygosity excess and mode-shift tests for all Lesser Kestrel populations. The Wilcoxon tests were carried using the TPM model (5% SMM, 95% IAM). Values in bold are indicative of a bottleneck (M<0.68 for the Garza-Williamson ratio and *P* < 0.05 for the Wilcoxon test)

| Code | Location | $N_E$ | M | $P$ (TPM) | Allele frequency distribution |
| --- | --- | --- | --- | --- | --- |
| ES | Spain | Not estimated | **0.43** | 0.083 | L-shaped |
| APU | Apulia | Not estimated | **0.45** | 0.148 | L-shaped |
| SIC | Sicily | 57.3<br>(29.8-327.7) | **0.39** | **0.007** | L-shaped |
| CRO | Croatia | 47.7<br>(21.6-82.8) | **0.41** | 0.216 | L-shaped |
| GIA | Ioannina | 26.6<br>(19.5-39.1) | **0.45** | 0.390 | L-shaped |
| LES | Agrinio | 88.4<br>(40.5-167.8) | **0.45** | 0.056 | L-shaped |
| CNG | Central-North<br>Greece | Not estimated | **0.45** | 0.078 | L-shaped |
| LIM | Limnos | 24.2<br>(14.5-55.8) | **0.42** | **0.004** | L-shaped |
| ISR | Israel | 54.9<br>(35.8-106.1) | **0.43** | **0.009** | L-shaped |
| MON | Mongolia | Not estimated | **0.42** | 0.201 | L-shaped |

## DISCUSSION

Across all sampled populations, STRUCTURE analysis suggested the presence of two major clusters concurring with the longitudinal distribution of the species; a ‘western’ (European) and an ‘eastern’ (Asian) cluster that are both represented in the population of Israel. This result reflects the proposed phylogeographic pattern of the species; based on the mitochondrial Cyt b region, European and Asian populations were found to be divergent whereas birds from Israel seem to cluster with both of them indicating a degree of genetic mixing (Wink *et al*., 2004). Interestingly, a comparison of plumage patterns of Lesser Kestrels unveiled substantial differences between individuals of European and Chinese descent whereas birds from Asian/Middle East populations (including Israel) exhibit an intermediate plumage pattern (Corso *et al*., 2016). In addition, this eastern cluster is highly represented in the eastern Greek population of Limnos Island (LIM) and extends gradually up to central Greece suggesting a population consisting of birds of eastern origin possibly from populations of Asia Minor. Future inclusion of samples from Western Anatolia (i.e. Turkey) and the Middle East could shed light on such hypotheses.

Despite the high private allelic richness, suggesting that gene flow could be restricted (at some level), our analysis showed low F_ST_ values among the core populations of Europe (Spain, Apulia region in Italy and Central Greece) which is in concordance with previous larger-scale studies (Alcaide *et al*., 2009). D_est_ on the other hand, generally showed higher values of differentiation among populations including the core ones (ES and APU). We found significant levels of differentiation among CE Mediterranean populations and a correlation of pairwise genetic and geographic distance. We should note that STRUCTURE may overestimate genetic structure in datasets characterized by such correlation (Frantz *et al*., 2009). Correlation between genetic and geographic distance has been previously reported for the species at both local (Ortego *et al*., 2008b) and large spatial scales (Alcaide *et al*., 2008a; Alcaide *et al*., 2008b) and can be driven by the distance-dependent dispersal exhibited by the species (Serrano & Tella, 2003; Serrano *et al*., 2003; Ortego *et al*., 2008b). Some long-distance dispersal events that could have facilitated gene flow among populations have been reported across the CE Mediterranean (Gustin, Mendi & Pedrelli, 2011; Bounas *et al*. 2016a). Nevertheless, some of these movements took place in the ’50s when the species showed a wider distribution. Actually, restricted dispersal in a fragmented range, coupled with high philopatry rates, along with a relatively short generation time of the species (average lifespan of 4-6 years; Newton & Olsen, 1990; Negro, 1997), might have contributed to a more frequent individual turnover in the breeding colonies thus allowing relatively quick changes in allele frequencies that led to the patterns of genetic differentiation observed in the region (Ortego *et al*., 2008a).

Our results for genetic bottlenecks seem to be contrasting, at first glance. *M* ratios suggest that the reported past population declines have indeed left genetic bottleneck signs in all populations throughout the species’ range while tests for heterozygosity excess suggested bottlenecks only in three populations. Detection of a bottleneck using *M* ratios but not heterozygosity excess is expected when a bottleneck is older, more severe, and/or the population has recovered (Williamson-Natesan, 2005). Thus Lesser Kestrel populations have at some point in the past undergone a severe, prolonged bottleneck, whereas in some areas (ISR, SIC, LIM) they seem to have experienced more recent, population declines. The Israeli population (ISR) has gone through a steep decline and is estimated to be less than 10% of the population prior to 1950 (Liven-Schulman *et al*., 2004). The geographic position of Israel population could explain the increased levels of diversity despite its small N_E_ as a result of the genetic admixture of immigrants from other European or Asian populations in the area. Recent bottleneck signs were also detected in two of the peripheral populations in the CE Mediterranean region (SIC, LIM), that were also differentiated and showed a relatively limited gene flow with other populations (Fig. 3) and small N_E_ but nevertheless similar diversity patterns. Despite its small N_E_, loannina (GIA) population exhibits high levels of diversity and gene flow with both core populations of CE Mediterranean (APU, CNG), a process that can alleviate the bottleneck effects. Since Ioannina is a known premigratory site for the species (Bounas *et al*., 2016b), these high levels of diversity could be explained by immigrants from other populations, that is individuals that visit the site during premigration and return to breed there. Besides, it has been suggested that non-breeding distributions can shape the genetic structure of populations (Szczys, Oswald & Arnold, 2017).

Finally, genetic drift could also play a role on the observed genetic patterns of populations in the region, since they exhibit small size and limited gene flow with other populations. This is particularly evident in the small Croatian population. This population consists of only 25 pairs and was recently discovered (Mikulic *et al*., 2013). Therefore we are ignorant regarding its history, that is if the population was recently founded or present historically but not detected. The population did not show any signs of bottleneck but there seems to be a loss of rare alleles as well as signs of allele fixation (Table 1, Supporting information, Figure S4). Despite that, there is no observed decrease in heterozygosity, which could be consistent with a scenario of a recently founded population that experienced the effects of genetic drift, since heterozygosity can be relatively insensitive to the loss of rare alleles due to drift (Allendorf, 1986; Allendorf, Aitken & Luikart, 2013).

### Conservation implications

According to our results, the genetic structure of the CE Mediterranean populations of the Lesser Kestrel appears to reflect to some extent the demographic decline that led to its range fragmentation. Therefore, we suggest the management of the whole CE Mediterranean population as a single conservation unit (ESU). However, the population could be divided into four different management units (MUs) since three peripheral populations (CRO, LIM, SIC) showed limited migration rates and different allele frequencies from all the other populations. Thus a mixed strategy that aims to preserve the high diversity of the core populations of Italy and Greece, while focusing on the recovery of bottlenecked peripheral populations (SIC, LIM), should be appropriate for the conservation of the species in the region.

Translocations of individuals from the core populations to the peripheral or newly founded colonies could be used as an effective conservation action (Morandini *et al*., 2017). Based solely on our results from the microsatellite analysis, the use of birds from the core populations seems to be an acceptable action since they were found to exhibit high diversity and low differentiation from all the others. However, the fact that our results were based solely on the analysis of selectively neutral loci prevents us from conclusions regarding the adaptive and evolutionary consequences of such action (Holderegger, Kamm & Gugerli, 2006). Thus any future translocation programmes should consider maximizing both the genetic and the adaptive similarity between populations.

## ACKNOWLEDGEMENTS

We are grateful to Action for Wildlife and ANIMA rehabilitation centers, the Lesser Kestrel Recovery project team (LIFE11 NAT/BG/360), J. Hernandez-Pliego and G. Giglio for helping with sample collection. D. Vavylis, G. Vakis and E. Toli assisted in the fieldwork. LIFE for the Lesser Kestrel project team (LIFE+ 11 NAT/GR/1011), and namely K. Vlachopoulos, S. Polymeros and A. Evangelidis greatly helped with sampling in Central Greece. N. Chakarov and M. Stubbe kindly provided samples from Mongolia and we are thankful for that. V. Saravia, P. Kordopatis and S. Marin provided valuable comments on a previous draft of this manuscript. We are grateful to J.A. Allen and two anonymous reviewers for their comments that greatly improved this manuscript. This study was financially supported by a scholarship awarded to A.B. by the A.G. Leventis Foundation.

